# Ignoring fossil age uncertainty leads to inaccurate topology and divergence times in time calibrated tree inference

**DOI:** 10.1101/2020.01.14.906107

**Authors:** Joëlle Barido-Sottani, Nina van Tiel, Melanie J. Hopkins, David F. Wright, Tanja Stadler, Rachel C. M. Warnock

## Abstract

Time calibrated trees are challenging to estimate for many extinct groups of species due to the incompleteness of the rock and fossil records. Additionally, the precise age of a sample is typically not known as it may have occurred at any time during the time interval spanned by the rock layer.

Bayesian phylogenetic approaches provide a coherent framework for incorporating multiple sources of evidence and uncertainty. In this study, we simulate datasets with characteristics typical of Palaeozoic marine invertebrates, in terms of character and taxon sampling. We use these datasets to examine the impact of different age handling methods on estimated topologies and divergence times obtained using the fossilized birth-death process. Our results reiterate the importance of modeling fossil age uncertainty, although we find that the overall impact of fossil age uncertainty depends on both fossil taxon sampling and character sampling. When character sampling is low, different approaches to handling fossil age uncertainty make little to no difference in the accuracy and precision of the results. However, when character sampling is high, sampling the fossil ages as part of the inference gives topology and divergence times estimates that are as good as those obtained by fixing ages to the truth, whereas fixing fossil ages to incorrect values results in higher error and lower coverage. Modeling fossil age uncertainty is thus critical, as fixing incorrect fossil ages will negate the benefits of improved fossil and character sampling.

## Introduction

Estimating phylogenetic relationships and divergence times among species are key components of piecing together evolutionary and geological history. Approaches to building time trees in paleobiology have traditionally involved estimating the topology and branch lengths scaled to time in separate, sequential analyses (Bapst and Hopkins, 2017). Bayesian phylogenetic models make it possible to estimate these parameters in combination. An advantage of this joint inference is that temporal evidence can be used to inform the tree topology, in combination with character data, and the posterior output will better reflect the uncertainty associated with the results (Ronquist et al., 2012).

Statistically coherent models for incorporating extinct species into time calibrated tree inference only recently became available. In particular, the fossilized birth-death (FBD) process provides a joint description of the diversification and fossil sampling processes (Stadler, 2010; Heath et al., 2014). Under this model, extinct dated samples are considered as part of the tree, therefore contributing temporal information, and their phylogenetic position can be recovered, either as terminal branches (tips) or ancestral to other samples (sampled ancestors). This modeling framework has created enormous potential for incorporating more paleontological data into divergence time analyses and we are only just beginning to explore the impact and existing limitations of this approach.

Analyses using the FBD process can be divided into two categories depending on the amount of data available. The first category uses topological constraints which assign fossils to specific clades (Heath et al., 2014; Gavryushkina et al., 2014). In these analyses the position of the fossils in the tree is thus not part of the inference. The second category are so-called “total-evidence” approaches, which use morphological data to place the fossils on the tree as part of the inference (Ronquist et al., 2012; Zhang et al., 2015; Gavryushkina et al., 2017). Total-evidence analyses better reflect the uncertainty associated with fossil placement than analyses that fix the position of fossils and thus may lead to more accurate results, particularly in clades where the fossil taxonomy is contested. This approach can also be applied to entirely extinct groups, for which only morphological and no molecular data are available (Slater, 2015; Wright, 2017b; Wright and Toom, 2017; Paterson et al., 2019).

Simulations play an important role in testing the limits of tree inference methods. Different taxonomic groups and time periods are associated with different issues that contribute to challenges inferring topology and time, and a growing number of studies have sought to explore the performance of phylogenetic inference under the FBD model in different scenarios. Several studies have focused on specific model violations, including the impact of non-uniform sampling among living taxa (Zhang et al., 2015; Matschiner, 2019), non-uniform sampling of fossil taxa over time (Heath et al., 2014; Zhang et al., 2015; O’Reilly and Donoghue, 2019) or across lineages (Heath et al., 2014; Matschiner et al., 2017), as well as the effect of ignoring sampled ancestors (Gavryushkina et al., 2014). A clear consensus that emerges from this work is that higher sampling rates of taxa and characters result in better estimates of time and (when co-estimated) topology, provided model violation is not extreme. In an extensive set of simulations, Luo et al. (2019) examined the performance of total-evidence inference under the FBD model. This work indicated that a large degree of uncertainty is anticipated to be associated with the placement of extinct samples, for which only morphology is available, and that fossil sampling may ultimately outweigh the significance of other issues encountered in dating analyses, including character sampling and among-lineage rate variation.

Barido-Sottani et al. (2019a) focused on one particular aspect of the fossil record, namely the uncertainty associated with the age assigned to each fossil sample. As the age of fossils is established in reference to the geological record, fossil samples are not dated to a single value but rather to an interval of time; this is referred to hereafter as the “age range” of the sample. This uncertainty can be handled in FBD analyses by sampling fossil ages as part of the inference (Drummond and Stadler, 2016), but many studies in the existing literature chose instead to fix fossil ages to a single value, usually the midpoint of the age range (e.g. Larabee et al., 2016) or an age sampled uniformly at random inside the range (e.g. Grimm et al., 2015). Barido-Sottani et al. (2019a) tested these different approaches of handling fossil age uncertainty in analyses using topological constraints to place fossils and found that fixing the fossil ages to incorrect values led to important errors in divergence times estimates. Here, we extend the Barido-Sottani et al. (2019a) study to time calibrated tree inference under the FBD model using morphological data only. Using simulated datasets, we explore the impact of character and taxon sampling, approaches to handling fossil age uncertainty, and clock model priors on estimates of topology and divergence times. We also compare our results to those obtained using temporally unconstrained (i.e. non-time calibrated) Bayesian tree inference. Finally, we apply the FBD model and several different methods for handing fossil age uncertainty to an empirical dataset that is typical of those available for Paleozoic marine invertebrates.

## Methods

### Simulated datasets

The design of our simulation study is broadly based on features that are typical for datasets of Paleozoic marine invertebrates. In order to select parameter values that would reflect the size and scale (in terms of taxon and character sampling, and time span) we first tallied 81 studies of Paleozoic invertebrate groups, which included trilobites (67%), brachiopods (18%) and crinoids (15%). The majority of these studies used species as operational taxonomic units (OTUs) (77%), while the rest coded genera (23%). The results are summarised in Fig. S1 and Appendix Table 1. The typical size of these datasets was 25–35 taxa, at both taxonomic levels (mean for species OTUs = 25, mean for genus OTUs = 32), with a maximum of 85. For 63 of these studies we were able to estimate approximate time spans in millions of years (Myr). Across all studies, the typical time span was 50 Myr, with 85% <75 Myr. However, there was a large difference between taxonomic scales: the mean total time span for studies using species OTUs was 37 Myr, while the mean total time span for studies using genus OTU studies was 88 Myr. Interestingly, no relationship was observed between the number of taxa in the study and the total time span. The average number of characters was 35 for both species and genus levels, with an average of 60% binary characters.

Based on these empirical data characteristics, we established two main parameter settings that determined the number of fossils sampled during simulation, one based on the average size of empirical datasets (referred to as *low sampling*) and the other based on a more optimistic sampling scenario (referred to as *high sampling*). We also explored the effect of morphological matrix length (30, 300 or 3 000 characters). For each set of parameter values we simulated 50 replicates.

### Simulation of phylogenies and fossil samples

Trees were simulated under a constant rate birth-death process with speciation rate *λ* = 0.06/Myr and extinction rate *µ* = 0.045/Myr, using the R package `TreeSim` (Stadler, 2011), based on estimates of these parameters from an empirical study of Paleozoic crinoids (Wright, 2017a). The process was allowed to run for 130 Myr, which approximates a clade evolving from the beginning of the Ordovician (485.4 Ma) to the end of the Devonian (358.9 Ma).

Fossils were sampled on the complete phylogeny following a Poisson process with constant rate *ψ*, using the R package `FossilSim` (Barido-Sottani et al., 2019b). We rejected phylogenies with less than *n*_min_ or more than *n*_max_ sampled fossils and phylogenies for which the fossils spanned less than *t*_span_ Myr. Values for *ψ, n*_min_, *n*_max_ and *t*_span_ depended on the fossil sampling setting (high versus low sampling) and are detailed in Table 1.

**Table 1:**
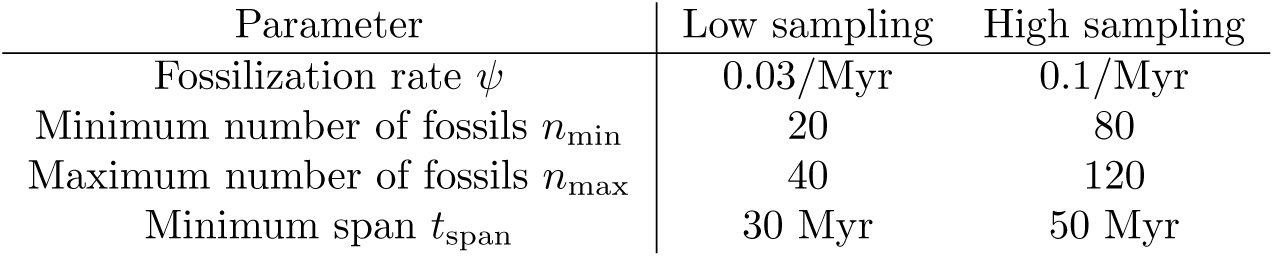
Parameters values for low and high fossil sampling settings.

| Parameter | Low sampling | High sampling |
| --- | --- | --- |
| Fossilization rate $\psi$ | 0.03/Myr | 0.1/Myr |
| Minimum number of fossils $n_{\min}$ | 20 | 80 |
| Maximum number of fossils $n_{\max}$ | 40 | 120 |
| Minimum span $t_{\text{span}}$ | 30 Myr | 50 Myr |

### Simulation of fossil age uncertainty

Fossil age uncertainty was simulated using the procedure described in Barido-Sottani et al. (2019a). Realistic age ranges for simulated data are based on empirical ranges of fossil crinoids obtained from the Paleobiology Database (PBDB) using the following parameters: time intervals = from Ordovician to Devonian, scientific name = Crinoidea. Each simulated fossil sample was assigned to an interval based on its true age. If a simulated fossil age could be assigned to multiple intervals, a single interval was selected at random by weighting all possible intervals by their frequency of appearance in the PBDB data. If no intervals appeared in the PBDB data for a simulated fossil age, a random interval containing the true age was drawn, with a length equal to the average length of all intervals in the PBDB data, i.e 12 Myr. Thus, the simulated interval for each fossil always included the correct age of the fossil.

### Simulation of morphological data

Binary characters were simulated for each fossil using the function `sim.char` from the R package `geiger` (Pennell et al., 2014). A strict clock model was used and the rate of character state change was set to 0.033/Myr, based on the rate obtained by Wright (2017a). For both fossil sampling settings, character matrices of length 30, 300 and 3 000 were simulated.

### Bayesian inference

Markov Chain Monte-Carlo (MCMC) inference using the FBD process is implemented in the Sampled Ancestors package (Gavryushkina et al., 2014) for the software BEAST2 (Bouckaert et al., 2014). We extended this package to be able to use a tree with no extant samples. This extension made no changes to the FBD model or to the likelihood function, and was done simply to allow for sampling fossil ages on a fully extinct tree. This package was used to perform Bayesian phylogenetic inference on the simulated datasets. The fossil ages were handled using five different methods, detailed here and illustrated in Figure 1.

**Figure 1:**
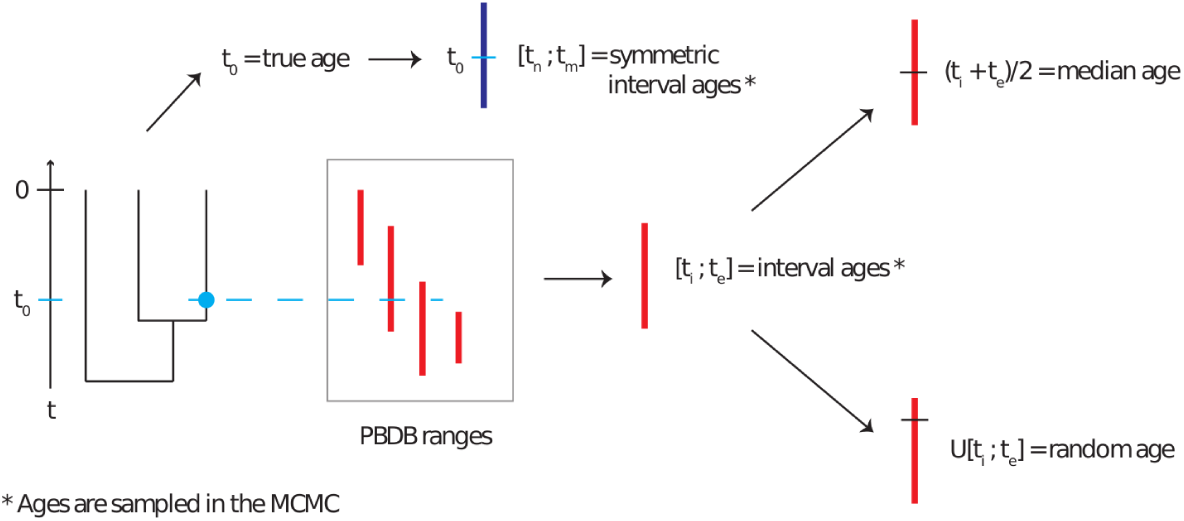
Representation of the age uncertainty simulation process, reproduced from Barido-Sottani et al. (2019a). Phylogenies with fossils are simulated according to a birth-death-fossilization process. The correct age of each fossil is used to draw an age interval for that fossil from the set obtained from PBDB. This age interval is then used as the basis for the median and random age assignment. A symmetric age interval is also drawn from the correct age.

- Correct ages: the fossil ages are fixed to the true ages as simulated.
- Interval ages: the fossil ages are not fixed, but are sampled along with the other parameters within the simulated age range.
- Median ages: the fossil ages are fixed to the midpoint of their simulated age range.
- Random ages: the fossil ages are fixed to an age sampled uniformly at random inside of their simulated age range.
- Symmetric interval ages: the fossil ages are not fixed, but are sampled along with the other parameters. Each fossil age is sampled within a symmetric interval around the true age of the fossil. The purpose of this setting was to evaluate the performance and accuracy in the situation where the midpoint of each prior interval was equal to the correct age.

The Lewis Mk model of morphological character evolution was used (Lewis, 2001). The strict clock model was used with three different priors on the clock rate: an unbounded uniform prior, a lognormal prior with median = 0.033/Myr, equal to the true rate (i.e. LogNormal(−3.4, 0.3)) and a lognormal prior with median = 1.220/Myr, different from the true rate (i.e. LogNormal(0.2, 1.25)). The inference was run for at least 100 000 000 iterations, or until convergence was considered satisfactory, and sampled every 10 000 steps. Convergence was assessed in the software Tracer v. 1.7 (Rambaut et al., 2018) and considered satisfactory if the effective sample sizes were more than 200. Datasets that did not converge after 120 hours were excluded; this only applied to a single replicate with high fossil sampling and character length 3 000.

Additionally, unconstrained Bayesian phylogenetic inferences were performed on the simulated datasets using the RevBayes framework (Höhna et al., 2016) without including any fossil age information. These inferences were performed on all simulated datasets for both low and high fossil sampling and, with character data simulated under the strict clock model. We used a uniform tree prior, an exponential branch length prior with mean = 1, and the Lewis Mk model. Convergence and MCMC diagnostics were assessed using identical guidelines as those described above.

### Assessing inference results

To assess the accuracy of inferred divergence times and FBD model parameters we measured the relative error of the median posterior estimates, where the relative error was defined as the difference between the true value and the estimated value, divided by the true value. The relative error was averaged over all replicates. We also calculated the coverage, i.e the proportion of analyses in which the true parameter value was included in the 95% highest posterior density (HPD) interval.

To assess the accuracy of inferred topologies we calculated the mean normalized Robinson-Foulds (RF) distance (Robinson and Foulds, 1981) between simulated trees and tree samples from the posterior distribution. The RF distance only depends on the topology of the trees. The normalized RF distance between two trees with *n* tips is computed by dividing the RF distance between these trees by the maximum possible RF distance between two trees with *n* tips, thus scaling the distances between 0 and 1.

The normalized RF distances were calculated using the `RF.dist` function from the R package `phangorn` (Schliep, 2010), and averaged over all the trees sampled during the MCMC (ignoring the first 10% of the samples as burn-in), and averaged over all replicates for each parameter combination.

All trees were unrooted prior to calculating the RF distance to facilitate comparison between the time constrained and unconstrained analyses.

### Empirical dataset

To explore the impact of different approaches to handling stratigraphic age uncertainty on empirical estimates of divergence times and topology, Bayesian phylogenetic inference was performed on a dataset of North American Devonian brachiopod species (Stigall Rode, 2005). The dataset is composed of 18 taxa and 36 characters, 80% of which are binary. As described in the methods (see also Fig. S1), matrices of this size are typical of those available for Paleozoic marine invertebrates. Fossil occurrences were assigned to geologic stages based on vetted occurrences in the Paleobiology Database and additional literature (Stigall Rode, 2005; Menning et al., 2006). Minimum and maximum ages for stage boundaries were assigned following the International Commission on Stratigraphy 2018 chart (www.stratigraphy.org). Species for which all specimens were recovered from the same geological stage were treated as a single OTU. Three species had specimens which were sampled from more than one stage and were treated as multiple OTUs, corresponding to one OTU for each stage. For these species, the species was constrained to be monophyletic and morphology was included for the oldest specimen only. This approach was taken in order to avoid having multiple specimens associated with the same morphology over long intervals of time, which would represent a strong violation of the Mk model. The analysis used the same model parameter-ization and priors as the simulated data. The clock rate prior was set to a lognormal distribution (LogNormal(−3.4, 0.3)). As the true ages of the fossils in this dataset are unknown, we limited our comparison to the median, random and interval ages in BEAST2, and the unconstrained analysis in RevBayes. To facilitate comparison between constrained and unconstrained topologies, unconstrained trees were rooted using *Xystostrophia umbraculum* as the outgroup taxon (Stigall Rode, 2005). However, we note that including the outgroup in the BEAST2 analyses had only a minor impact on the estimated ingroup topology or divergence times (results in supplement).

## Results

### Simulated datasets

#### Impact of the clock rate prior

The use of different clock rate priors had a minor impact on the results. For most parameters, including divergence times, estimates of coverage and relative error were similar or even identical for different clock priors, particularly when character and fossil sampling were both high (Figures 2,3). The use of different priors on the clock rate also had little impact on the topological accuracy based on RF distances (Figure 4). The largest impact was observed on the clock rate itself. When character and fossil sampling were low, there was less signal in the data to inform this parameter. Running the analysis with an unbounded uniform prior (i.e. from 0 to ∞) on the clock rate in the low sampling scenario produced rate estimates of ≈ 1^307^ (i.e. the numerical upper limit of the software) in approximately 20% of replicates, as the posterior followed the prior. Thus, we excluded this condition from Figure 2 (lower panel).

**Figure 2:**
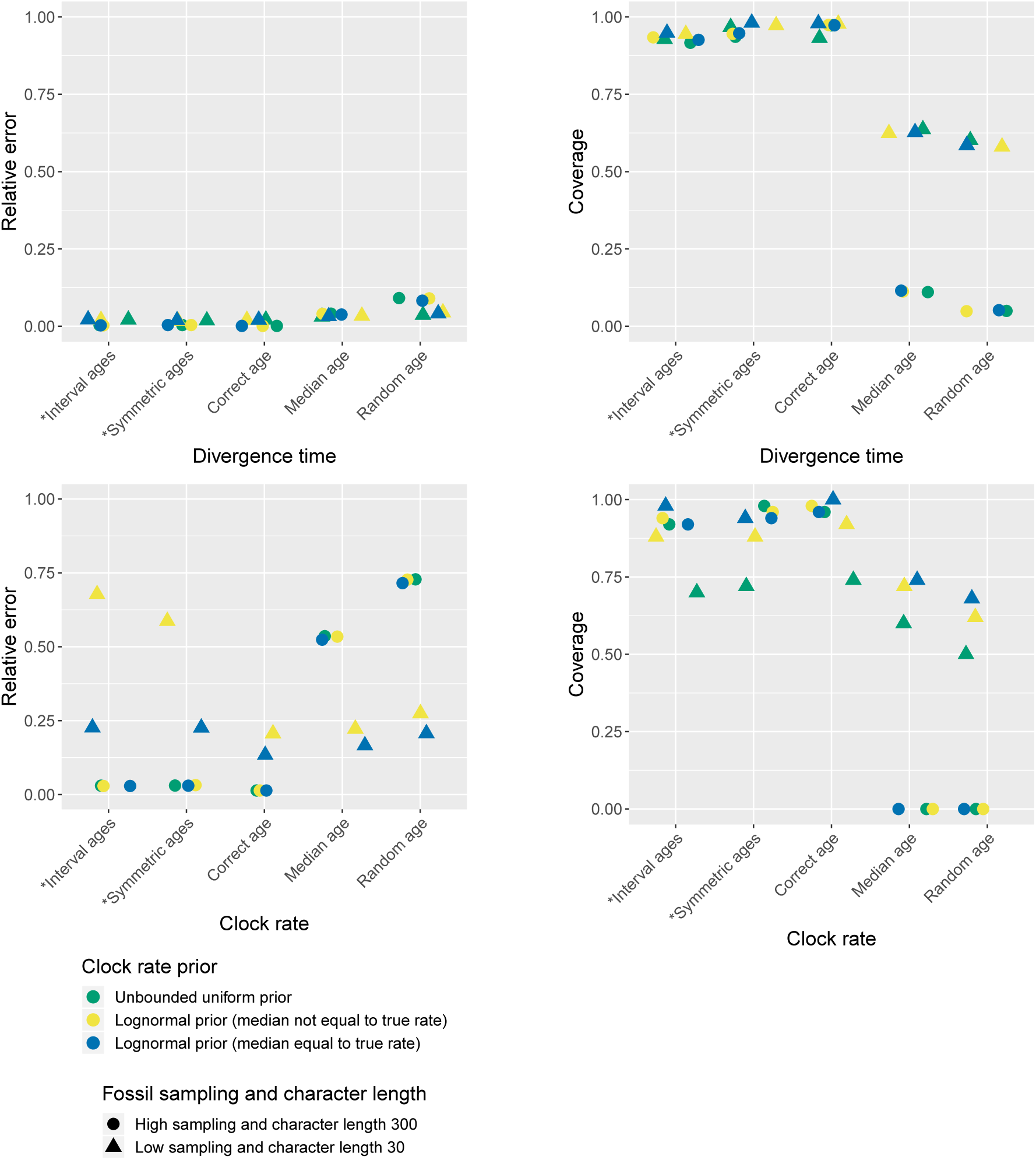
Impact of the clock prior on divergence times and clock rate. Average relative error of median posterior estimates and 95% HPD coverage are shown for different clock rate priors, different age handling methods, and different fossil sampling settings. Ages sampled as part of the MCMC are marked by (*).

**Figure 3:**
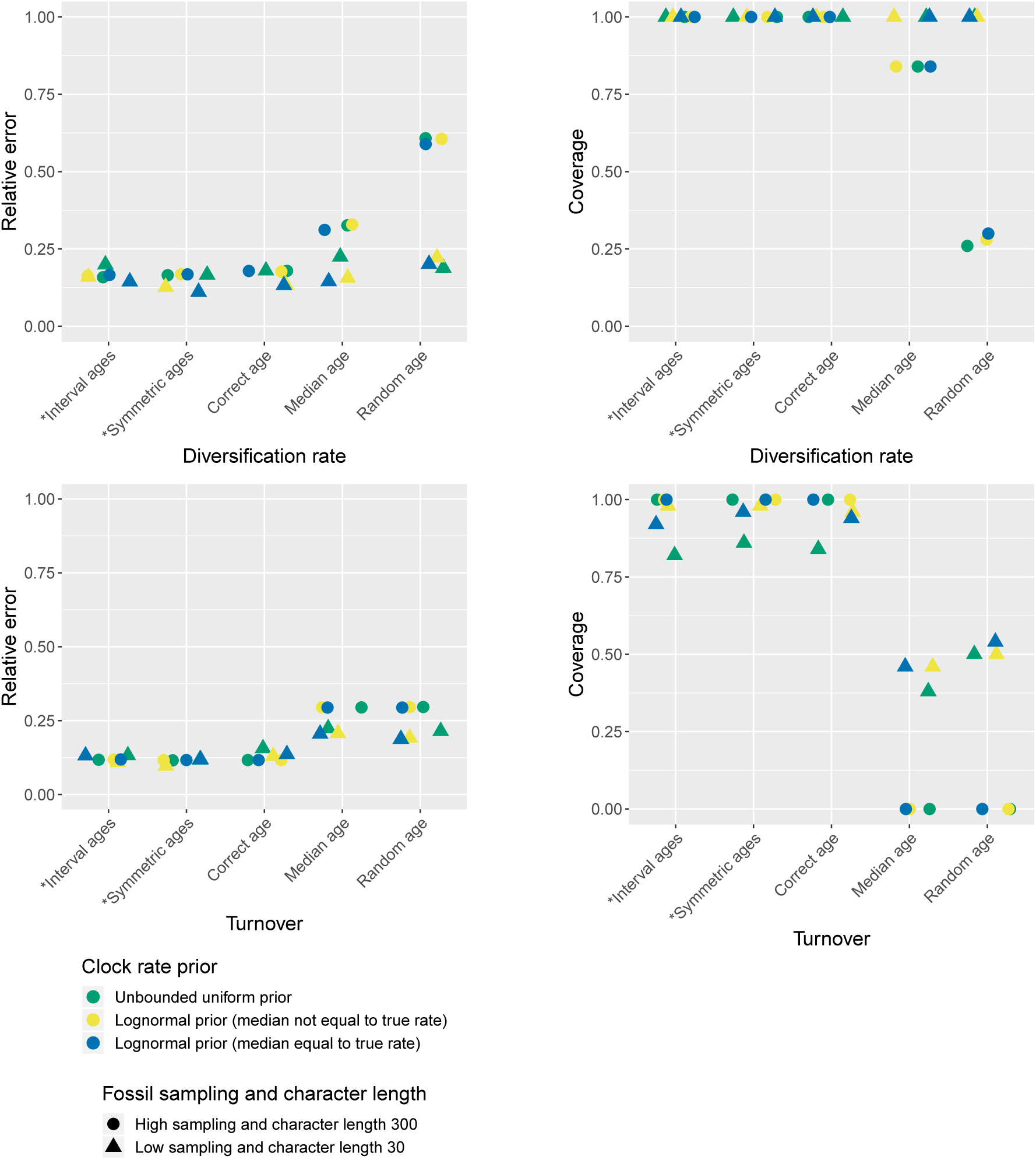
Impact of the clock prior on diversification and turnover. Average relative error of median posterior estimates and 95% HPD coverage are shown for different clock rate priors, different age handling methods, and different fossil sampling settings. Ages sampled as part of the MCMC are marked by (*).

**Figure 4:**
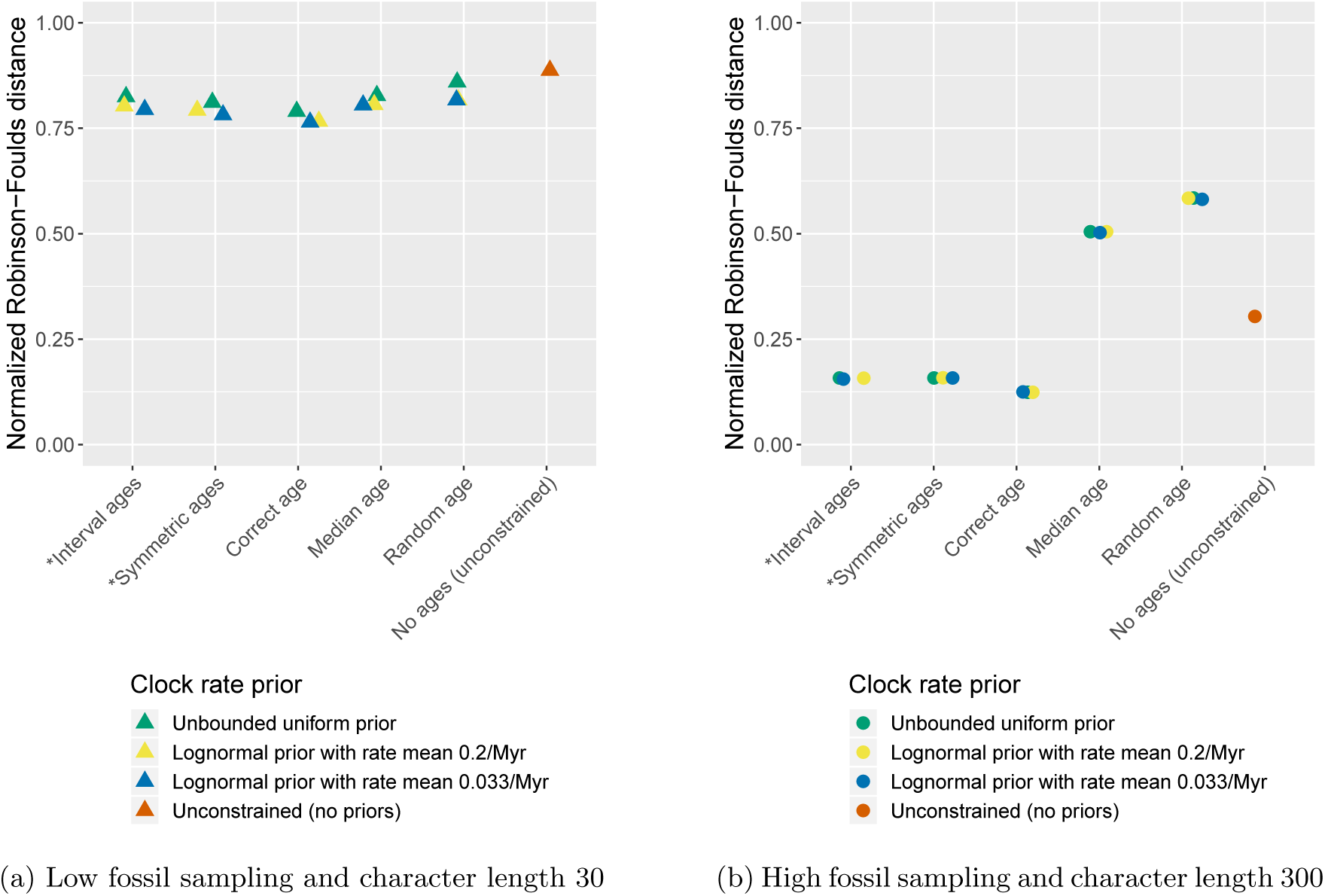
Impact of the clock prior on topology. Average normalized Robinson-Foulds distance between the true simulated tree and trees inferred during MCMC across all replicates are shown for different clock rate priors, different age handling methods, and different fossil sampling settings (low (a) versus high (b)). Ages sampled as part of the MCMC are marked by (*).

#### The combined effects of stratigraphic age uncertainty, fossil sampling, and character sampling

Figures 5–7 present the results obtained under different character and fossil sampling settings when running the analysis with a prior on the clock rate set to a lognormal distribution with a median that differs from the true rate. The accuracy of inferred divergence times, in terms of coverage and relative error, show similar behavior across fossil and character sampling settings (see Figure 5). In particular, we obtained high accuracy (i.e. high coverage and low relative error) when the fossil ages were fixed to the correct ages or sampled from within the known interval of uncertainty as part of the MCMC, irrespective of fossil or character sampling. In contrast, we obtained low accuracy when the ages were fixed to incorrect (median or random) ages, but the extent to which the results were worse depended on both fossil and character sampling. In the case of fixed incorrect ages, increased fossil and character sampling decreased the accuracy of divergence time estimates. A similar trend is observed for the diversification and turnover parameters (see Figure 6). The clock rate parameter showed the same trends for coverage (i.e. higher fossil and character sampling lead to lower coverage with median or random fossil ages), but a different trend was recovered for relative error (see Figure 5). Specifically, when fossil and character sampling were low, relative error was higher when fossil ages were co-estimated compared to when the ages were fixed to either the correct or incorrect ages. However, coverage was consistently lower with incorrect fossil ages.

**Figure 5:**
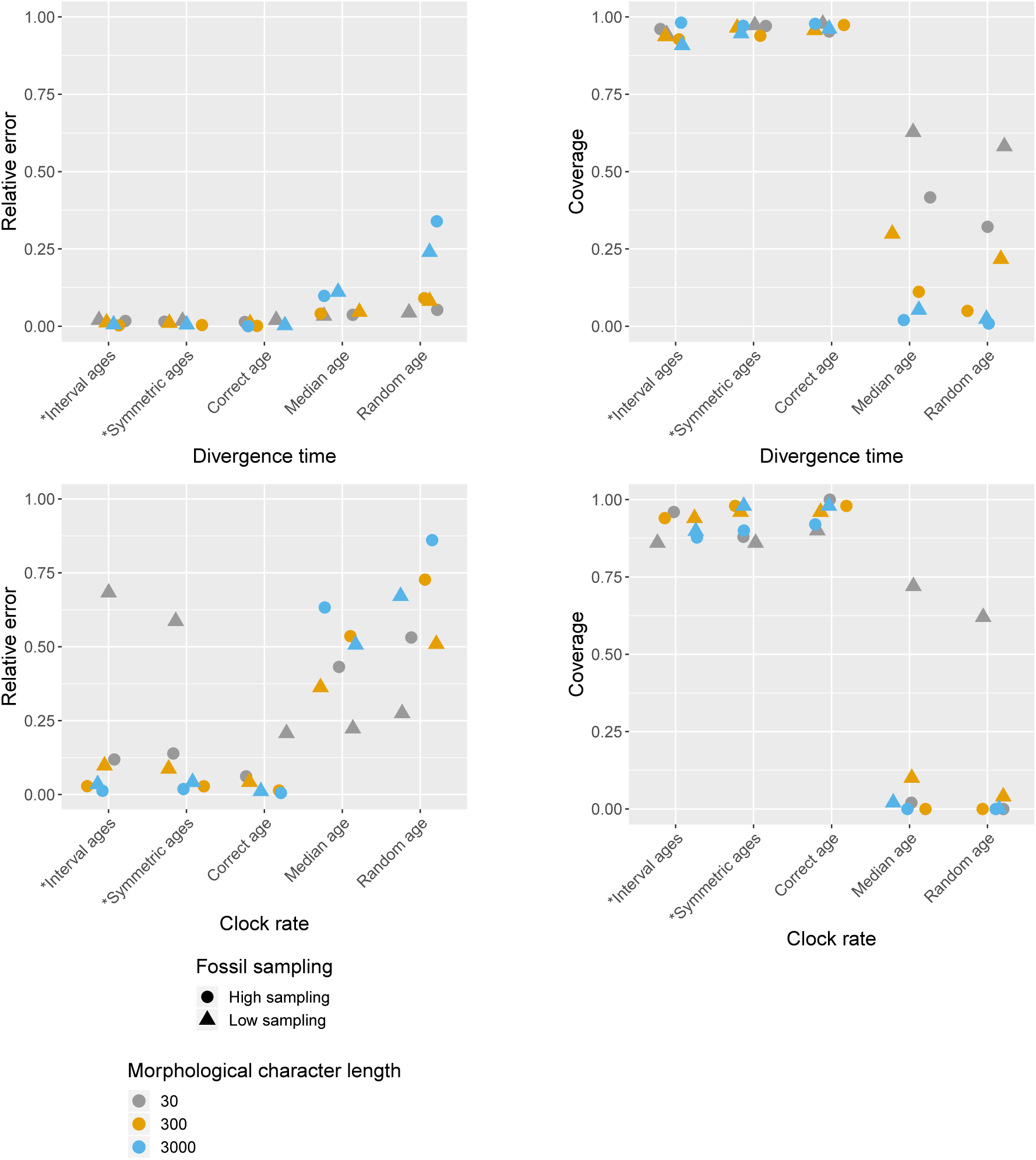
Impact of character and fossil sampling on divergence times and clock rate. Average relative error of median posterior estimates and 95% HPD coverage are shown for different age handling methods, different character sampling and different fossil sampling settings. Ages sampled as part of the MCMC are marked by (*).

**Figure 6:**
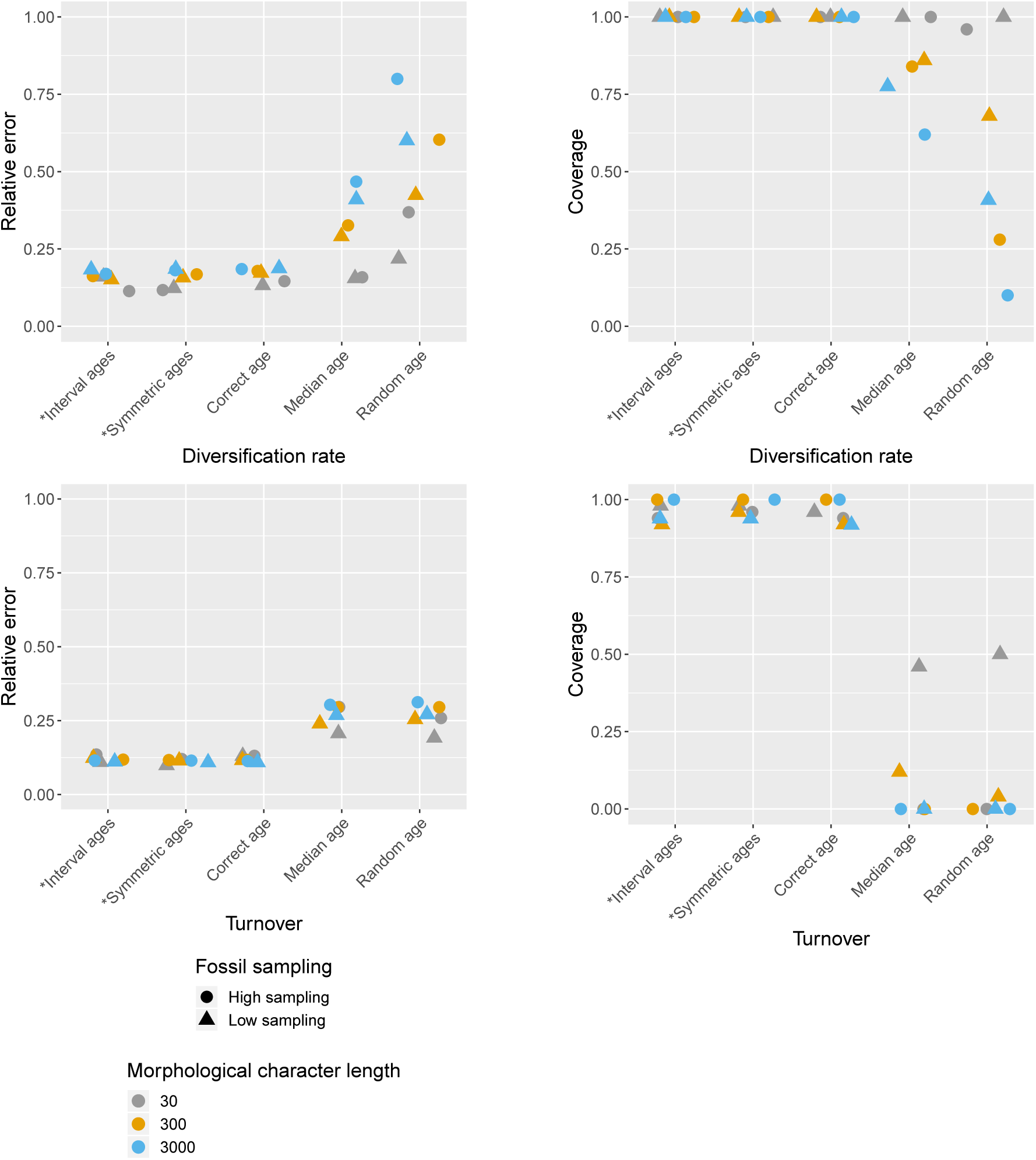
Impact of character and fossil sampling on diversification and turnover. Average relative error of median posterior estimates and 95% HPD coverage are shown for different age handling methods, different character sampling and different fossil sampling settings. Ages sampled as part of the MCMC are marked by (*).

**Figure 7:**
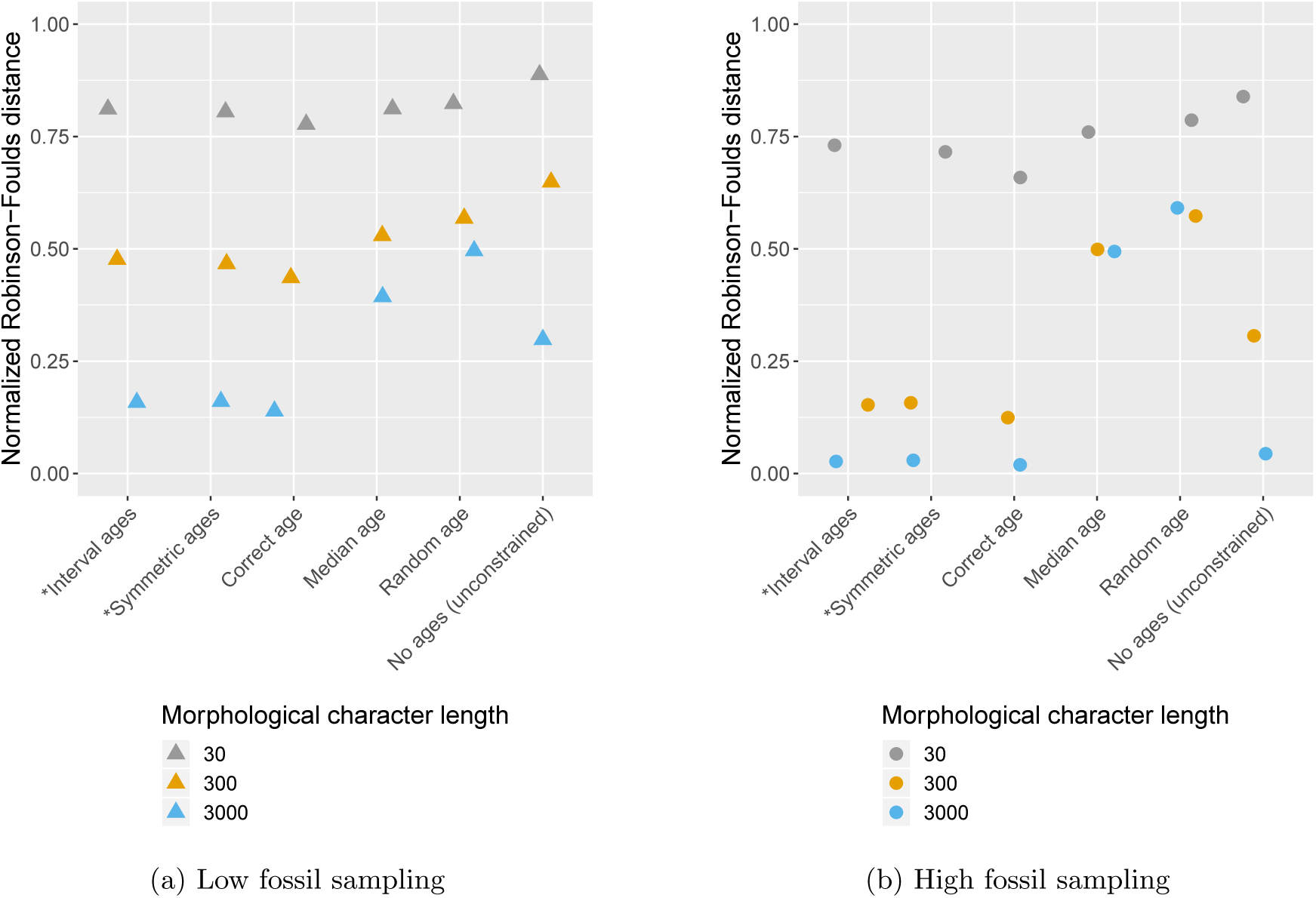
Impact of character and fossil sampling on topology. Average normalized Robinson-Foulds distance between the true simulated tree and trees inferred during MCMC across all replicates are shown for shown for different age handling methods, different character sampling and different fossil sampling settings (low (a) versus high (b)). Ages sampled as part of the MCMC are marked by (*).

The accuracy of inferred trees follow a pattern which is similar overall to the divergence times estimates, across fossil age handling approaches and fossil sampling settings. However, character sampling had a large impact on the magnitude of the differences observed under different age handling and fossil sampling scenarios (Figure 7). In particular, when character sampling was low (*n* = 30) the inferred trees were relatively far from the true tree, as measured by RF distance, irrespective of fossil age handling approach or fossil sampling parameters. Overall, higher character and fossil sampling both led to increased accuracy (i.e. lower RF distances) across all scenarios, with the best estimates obtained when both character and fossil sampling were high (Figure 7). The positive effects of increased fossil or character sampling were also greater when fossil ages were fixed to the truth or co-estimated, while estimates obtained when fossil ages were fixed to median or random ages remained inaccurate even with high sampling. When fossil and character sampling were both high, using the correct fossil ages or estimating the ages performed much better than using incorrect fossil ages.

Differences in accuracy between time calibrated and unconstrained tree inferences were also linked to variation in character sampling (Figure 4). For low or intermediate character sampling (*n* = 30 or 300) combined with low fossil sampling, or for low character sampling (*n* = 30) combined with high fossil sampling, the FBD inference outperformed the unconstrained inference, irrespective of the fossil age handling method. In contrast, for increased character or fossil sampling (*n* = 3000 combined with low fossil sampling and *n* = 300 or 3000 combined with high fossil sampling), the unconstrained inference outperformed the FBD model when fossil ages were fixed to incorrect ages. The FBD model outperformed the unconstrained inference under intermediate sampling scenarios (*n* = 3000 combined with low fossil sampling and *n* = 300 combined with high fossil sampling) when fossil ages were fixed to the correct ages or co-estimated. When fossil and character sampling were both high the results obtained using both constrained and unconstrained analyses converged on the true tree, provided fossil ages were fixed to correct ages or co-estimated.

The results are summarized in Table 2. Overall, these results indicate that increasing the amount of data does not compensate for the errors introduced by fixing fossil ages to incorrect values. On the contrary, these errors have a much larger impact when using larger datasets, to the point that discarding the fossil ages entirely leads to better estimates of the topology than using incorrect fixed ages.

**Table 2:**
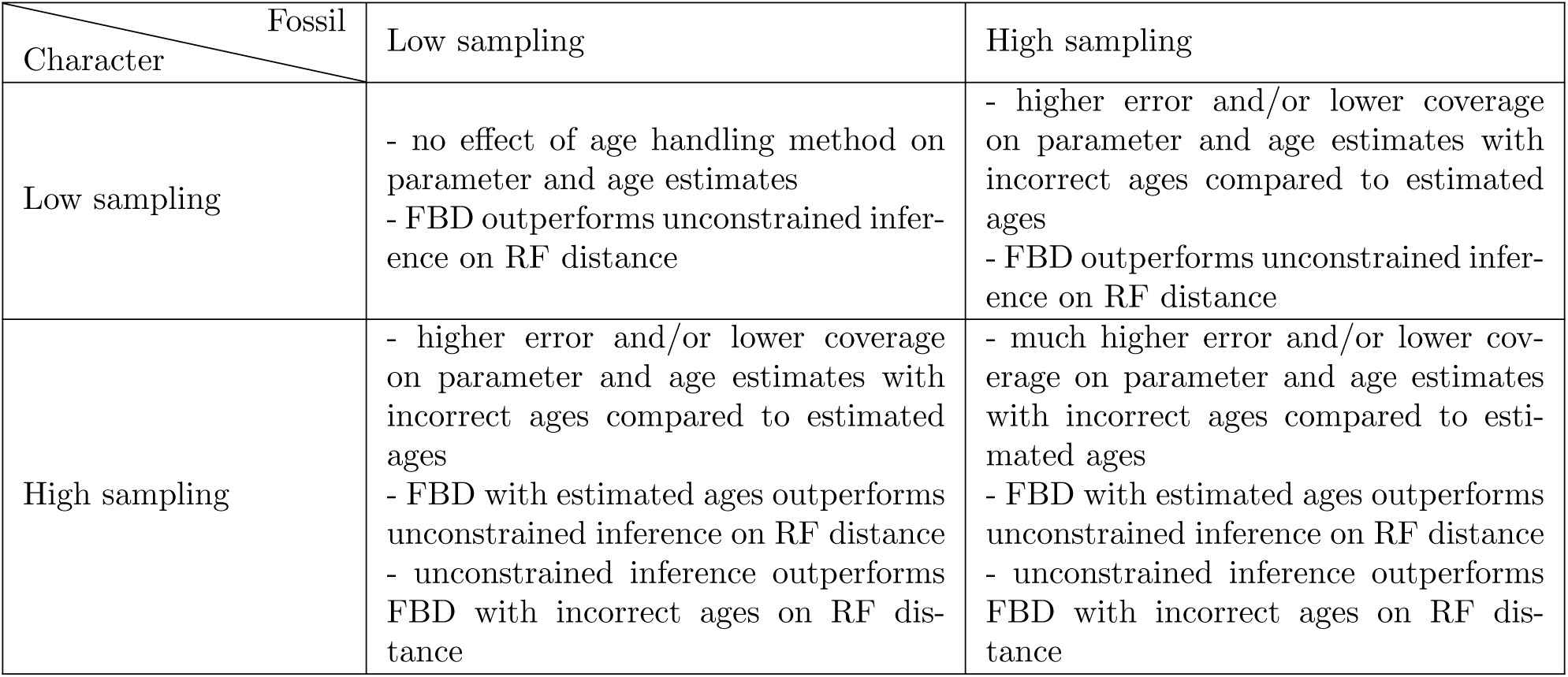
Impact of fossil and character sampling on the estimates obtained using the FBD model with different age handling methods versus unconstrained (i.e. non-clock) inference.

#### Empirical dataset

The Maximum Clade Credibility (MCC) trees obtained with interval ages, median ages, random ages or unconstrained analysis using our empirical brachiopod dataset are shown in Figure 8. The parameter estimates obtained under different age handling methods are shown in Figure S2. All OTUs belonging to the same species were constrained to be monophyletic. However, the posterior support for these nodes may be lower than 1.0, as TreeAnnotator considers the clade (A1(A2)), where A1 is a sampled ancestor of A2, to be different from the clade (A1,A2) and thus counts them separately when calculating their support.

**Figure 8:**
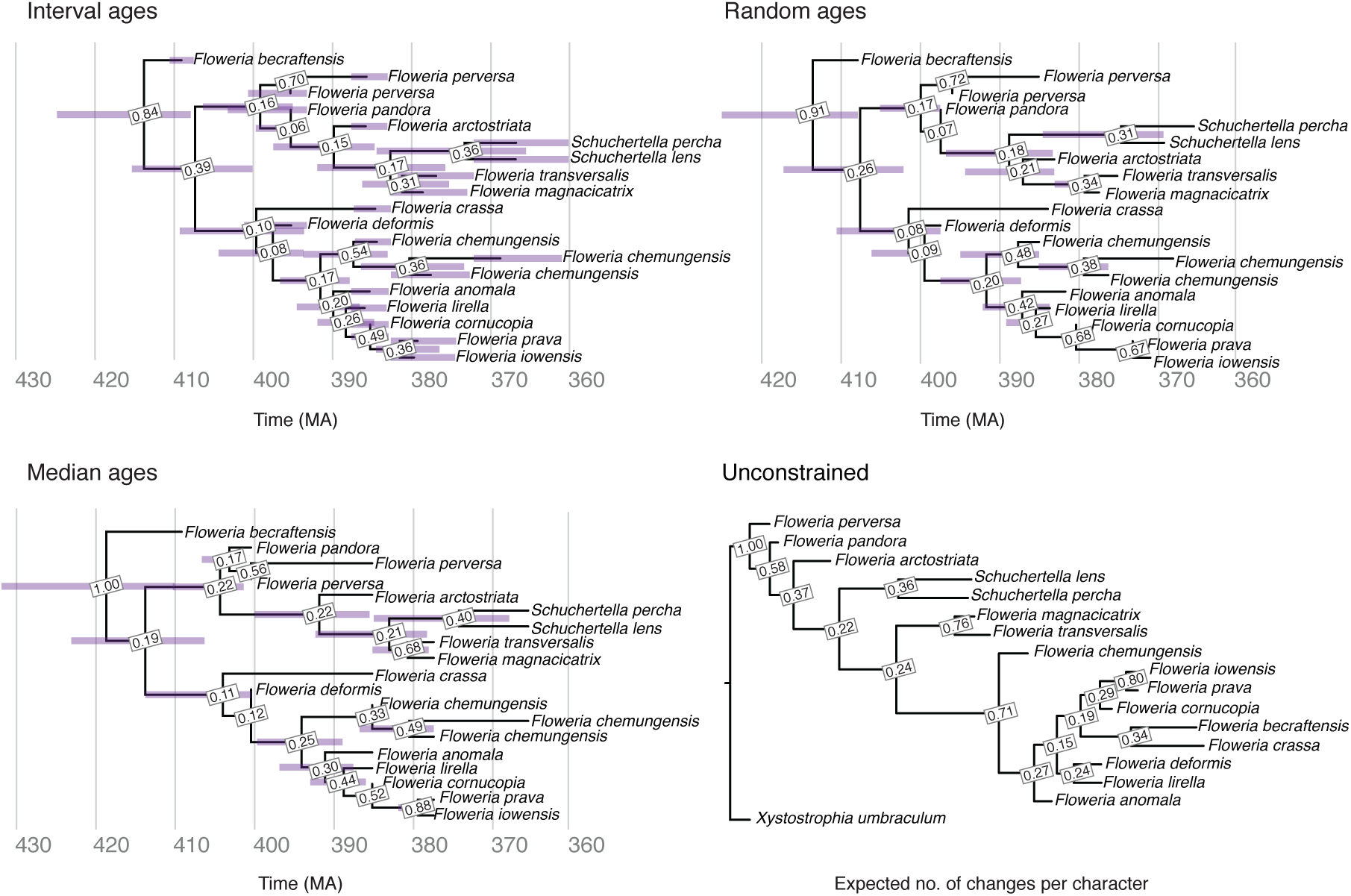
Brachiopod MCC trees obtained using the FBD analysis with interval ages, median ages and random ages, and unconstrained analysis. Posterior support is shown for each node for all trees. Error bars on the FBD trees show the 95% HPD interval for the age of each node, as well as the age of each fossil in the tree with interval ages.

The MCC trees obtained with the three methods for handling fossil ages are all almost identical in terms of their topology, with the exception of the placement of *Floweria arctostriata* in the random ages tree, and the node support is consistent across all three analyses. The median root ages are slightly different, with the median root age for the interval ages analysis the youngest, but only by a few million years (Figures 8 and S2). The MCC for the unconstrained analysis supports some of the same sister taxa with similar support values, and larger subclades are broadly consistent with several exceptions for specific taxa. Particularly notable is the derived placement of *Floweria becraftensis* in the unconstrained analysis. This species is among the oldest of the clade, and when fossil ages are included in the analysis, it is placed in a more basal position.

## Discussion

The FBD model can be used to estimate time calibrated trees under a range of scenarios. Our goal was to examine the impact of stratigraphic age uncertainty in FBD model analyses for datasets that are characteristic of fully extinct clades, such as Paleozoic marine invertebrate groups. Our survey of empirical data confirms that taxonomic groups from this time period typically have a small number of both taxa and phylogenetic characters. The age uncertainty associated with fossil samples from this time period is also relatively high (12 Myr on average). Our results demonstrate the importance of incorporating stratigraphic age uncertainty into phylogenetic dating analyses on these datasets, rather than the popular practice of fixing fossil ages to a value from within the known interval of uncertainty, e.g. using the mean or a random value (Figures 2,3,5,6).

Our results build on the findings of our previous work, where we showed that fixing fossil ages to incorrect values can lead to inaccurate estimates of divergence times under the FBD model when using topological constraints to place the fossils (Barido-Sottani et al., 2019a). In this previous study, we focused on a scenario where the aim was to estimate divergence times among extant species using molecular data. No character data was available for fossil samples but it was assumed that strong prior information was available to constrain the topology. Here, we assumed that the phylogenetic position of fossil samples was unknown and used morphological data to co-estimate topology along with divergence times. The results of our simulations show that in addition to recovering inaccurate divergences times, mishandling fossil age uncertainty can also result in the wrong tree (Figures 4,7).

We did not examine the impact of non-uniform fossil recovery, though this is known to decrease performance of the FBD model if unaccounted for (Heath et al., 2014; O’Reilly and Donoghue, 2019; Luo et al., 2019). Overall, our simulated datasets were designed to represent a best-case scenario for a fully extinct Paleozoic clade. We anticipate that additional, unaccounted-for model violations, such as non-uniform fossil recovery, would increase the errors in topology and divergence times estimates reported in this study.

Similarly, we did not examine the impact of morphological model violations such as rate heterogeneity among characters or the effects of non-uniform missing character data. A recent study suggested that even large deviations from the true model may have limited impact on divergence times estimates using total-evidence dating under the uniform tree model (Klopfstein et al., 2019). However, none of their simulation scenarios excluded molecular data and thus these findings may not be applicable to fully extinct clades. That said, the overall number of phylogenetic characters may be more of a concern for extinct clades, given the large degree of uncertainty associated with small matrices.

Small character matrices can be due to low taxon sampling, low character sampling, or both. The effect of both has been examined in previous studies. For example, simulations focused on unconstrained (i.e. non time calibrated) Bayesian inference have shown that small morphological matrices (e.g. 100 characters or less) will result in highly uncertain trees (Puttick et al., 2017; O’Reilly and Donoghue, 2017). Taxon sampling is also important for phylogenetic accuracy in unconstrained tree inference (Heath et al., 2008). Several simulation studies have demonstrated the importance of having sufficient fossil sampling in order to recover reliable estimates of divergence times using the FBD model (Heath et al., 2014; O’Reilly and Donoghue, 2019). Luo et al. (2019) examined the combined effects of fossil and character sampling on total-evidence estimates of time and topology, including a scenario that used morphological data only. Similar to our findings, their results show that increasing both the number of fossil samples and morphological characters leads to better estimates of time and topology, in terms of accuracy and precision. They also compared the use of fixed versus co-estimated fossil ages, where the age of fossils were fixed to the truth or ages were estimated from within the known interval of uncertainty. They found no strong differences in the estimated node ages when co-estimating fossil ages, which is coherent with our simulation scenarios, in which we observe very little difference in accuracy when comparing true versus co-estimated fossil ages.

Our simulations also show that the inclusion of fossil age information can improve the inferred topology regardless of the size of the matrix, if fossil age uncertainty is handled appropriately (Figures 4,7). On the other hand, excluding age information is preferable to using incorrect fossil ages even when using large morphological matrices. Thus, stratigraphic age uncertainty must be taken into account in order to fully benefit from the inclusion of fossil sampling times in the analysis.

Assuming that fossil age uncertainty is handled appropriately, our results indicate that the priority for improving topology and divergence times should be to increase matrix size. However, some clades are naturally small or rare. For these clades, even with complete taxon sampling, the size of the dataset will remain small. The best course of action then may be to increase the taxonomic scope of the study and to sample more broadly. In the case of fossil clades, small numbers of characters may reflect the paucity of morphological trait data available from some groups whose record is characterized by exoskeletal or shell elements exhibiting minimal morphological variation. However, small matrix size might also reflect the historical circumstances in which these data were generated: many matrices surveyed (Appendix S1) were constructed for parsimony analysis where the focus was on the selection of phylogenetically informative characters and not necessarily intended to represent an exhaustive survey of the preserved variation. Moreover, some workers report they excluded a subset of characters from consideration because of *a priori* concerns about homoplasy (Guensburg and Sprinkle, 2003), and therefore only sample characters they considered relevant or taxonomically significant. In this regard, it is conceivable that many published matrices may be expanded by a resurvey of the taxa of interest.

In addition, continuous trait data could provide an additional source of morphological information to complement matrices of discrete characters. Models of continuous trait evolution can be used to infer topology (Parins-Fukuchi, 2017) and divergence times (Alvarez-Carretero et al., 2019). Continuous data has also been shown to capture higher phylogenetic signal compared to discrete characters and can result in more accurate trees (Parins-Fukuchi, 2018). It is worth noting that >30% of the characters in our empirical example using brachiopods are continuous characters broken down into discrete states. If the process of discretization results in a loss of phylogenetic information, then tree inference and divergence time estimation could potentially be improved by modelling discrete and continuous characters separately.

We note that previous simulations examining the performance of both unconstrained or time calibrated Bayesian phylogenetic inference tend to use a minimum of 100 characters, which is > 3 times the size of datasets available for many fossil invertebrate groups. Matrices of only 20-30 characters, which are widely used in the literature, may contain too much uncertainty for other methodological choices to matter. Thus, we must be realistic about the degree of uncertainty expected when the number of phylogenetic characters sampled is low. All approaches to constructing summary trees are problematic when there is a lot of uncertainty in the posterior and all summary trees should be interpreted with caution (O’Reilly and Donoghue, 2017). In conclusion, we show that as more phylogenetically informative data become available, fixing the fossil ages to incorrect values can lead to important errors. Sampling fossil ages as part of the inference recovers estimates similar to those obtained when fixing the ages to the correct values. Consequently, we recommend incorporating stratigraphic age uncertainty when conducting analyses using the FBD process.

## Data availability

The extended SA package is available on GitHub https://github.com/CompEvol/sampled-ancestors and via the BEAST2 package manager. The R scripts used to simulate and analyse the data, as well as the XML files used to run BEAST2, will be made available on GitHub.

## Author contributions

JBS and NvT implemented the study, ran the simulations and analyzed the results. MJH and DFW assembled the empirical datasets and analyzed the empirical results. TS provided feedback on the study design and results. RCMW designed the study and interpreted the results. All authors contributed to writing the manuscript.

## Acknowledgements

This is Paleobiology Database official publication XXX. J.B.S was supported by funds from the National Science Foundation (US), grant DBI-1759909. D.F.W acknowledges support from the Gerstner Scholars Fellowship and the Gerstner Family Foundation, the Lerner-Gray Fund for Marine Research, and the Richard Gilder Graduate School, American Museum of Natural History, as well as a Norman Newell Early Career Grant from the Paleontological Society. R.C.M.W. was funded by the ETH Zürich Postdoctoral Fellowship and Marie Curie Actions for People COFUND programme.

## References

Sandra Alvarez-Carretero, Anjali Goswami, Ziheng Yang, and Mario Dos Reis. Bayesian estimation of species divergence times using correlated quantitative characters. Systematic Biology, page syz015, 2019. doi: 10.1093/sysbio/syz015.

David W. Bapst and M.J. Hopkins. Comparing cal3 and other a posteriori time-scaling approaches in a case study with the pterocephaliid trilobites. Paleobiology, 2017.

Jöelle Barido-Sottani, Gabriel Aguirre-Fernández, Melanie Hopkins Hopkins, Tanja Stadler, and Rachel Warnock. Ignoring stratigraphic age uncertainty leads to erroneous estimates of species divergence times under the fossilized birth-death process. Proceedings of the Royal Society of London B: Biological Sciences, 2019a.

Jöelle Barido-Sottani, Walker Pett, Joseph E O’Reilly, and Rachel CM Warnock. Fossilsim: an r package for simulating fossil occurrence data under mechanistic models of preservation and recovery. Methods in Ecology and Evolution, 2019b.

Remco Bouckaert, Joseph Heled, Denise Kühnert, Tim Vaughan, Chieh-Hsi Wu, Dong Xie, Marc A. Suchard, Andrew Rambaut, and Alexei J. Drummond. Beast 2: A software platform for bayesian evolutionary analysis. PLOS Computational Biology, 10(4):1–6, 2014. doi: 10.1371/journal.pcbi.1003537. URL https://doi.org/10.1371/journal.pcbi.1003537.

Alexei J Drummond and Tanja Stadler. Bayesian phylogenetic estimation of fossil ages. Philosophical Transactions of the Royal Society B: Biological Sciences, 371(1699):20150129, 2016.

Alexandra Gavryushkina, David Welch, Tanja Stadler, and Alexei J. Drummond. Bayesian inference of sampled ancestor trees for epidemiology and fossil calibration. PLoS computational biology, 10(12):e1003919, 2014. ISSN 15537358. doi: 10.1371/journal.pcbi.1003919. URL http://journals.plos.org/ploscompbiol/article?id=10.1371/journal.pcbi.1003919.

Alexandra Gavryushkina, Tracy A Heath, Daniel T Ksepka, Tanja Stadler, David Welch, and Alexei J Drummond. Bayesian total-evidence dating reveals the recent crown radiation of penguins. Systematic Biology, 66(1):57–73, 2017.

G. W. Grimm, P. Kapli, B. Bomfleur, S. McLoughlin, and S. S. Renner. Using More Than the Oldest Fossils: Dating Osmundaceae with Three Bayesian Clock Approaches. Systematic Biology, 64 (3):396–405, may 2015. ISSN 1063-5157. doi: 10.1093/sysbio/syu108. URL https://academic.oup.com/sysbio/article-lookup/doi/10.1093/sysbio/syu108.

T.E. Guensburg and J. Sprinkle. The oldest known crinoids (early ordovician, utah) and a new crinoid plate homology system. Bulletins of American Paleontology, 364, 2003.

Tracy A Heath, Shannon M Hedtke, and David M Hillis. Taxon sampling and the accuracy of phylogenetic analyses. Journal of Systematics and Evolution, 46(3):239–257, 2008.

Tracy A. Heath, John P. Huelsenbeck, and Tanja Stadler. The fossilized birth-death process for coherent calibration of divergence-time estimates. Proceedings of the National Academy of Sciences of the United States of America, 111(29):E2957–66, jul 2014. ISSN 1091-6490. doi: 10.1073/pnas.1319091111. URL http://www.ncbi.nlm.nih.gov/pubmed/25009181 http://www.pnas.org/content/111/29/E2957.abstract.html?etoc http://www.pnas.org/cgi/doi/10.1073/pnas.1319091111 http://www.pubmedcentral.nih.gov/articlerender.fcgi?artid=PMC4115571.

Sebastian Höhna, Michael J. Landis, Tracy A. Heath, Bastien Boussau, Nicolas Lartillot, Brian R. Moore, John P. Huelsenbeck, and Fredrik Ronquist. RevBayes: Bayesian Phylogenetic Inference Using Graphical Models and an Interactive Model-Specification Language. Systematic Biology, 65(4):726–736, 05 2016. ISSN 1063-5157. doi: 10.1093/sysbio/syw021. URL https://doi.org/10.1093/sysbio/syw021.

Seraina Klopfstein, Remo Ryser, Mario Corio, and Tamara Spasejovic. Mismatch of the morphology model is mostly unproblematic in total-evidence dating: insights from an extensive simulation study. bioRxiv, page 679084, 2019.

Fredrick J. Larabee, Brian L. Fisher, Chris A. Schmidt, Pável Matos-Maraví, Milan Janda, and Andrew V. Suarez. Molecular phylogenetics and diversification of trap-jaw ants in the genera Anochetus and Odontomachus (Hymenoptera: Formicidae). Molecular Phylogenetics and Evolution, 103:143–154, oct 2016. ISSN 1055-7903. doi: 10.1016/J.YMPEV.2016.07.024. URL https://www.sciencedirect.com/science/article/pii/S1055790316301804{#}s0010.

Paul O. Lewis. A Likelihood Approach to Estimating Phylogeny from Discrete Morphological Character Data. Systematic Biology, 50(6):913–925, 11 2001. ISSN 1063-5157. doi: 10.1080/106351501753462876. URL https://doi.org/10.1080/106351501753462876.

Arong Luo, David A Duchêne, Chi Zhang, Chao-Dong Zhu, and Simon Ho. A Simulation-Based Evaluation of Tip-Dating Under the Fossilized Birth-Death Process. Systematic Biology, 2019. ISSN 1063-5157. doi: 10.1093/sysbio/syz038.

Michael Matschiner. Selective sampling of species and fossils influences age estimates of the fossilized birth-death model. Frontiers, 2019.

Michael Matschiner, Zuzana Musilová, Julia MI Barth, Zuzana Starostová, Walter Salzburger, Mike Steel, and Remco Bouckaert. Bayesian phylogenetic estimation of clade ages supports trans-atlantic dispersal of cichlid fishes. Systematic Biology, 66(1):3–22, 2017.

Manfred Menning, AS Alekseev, BI Chuvashov, VI Davydov, F-X Devuyst, HC Forke, TA Grunt, Luc Hance, PH Heckel, NG Izokh, et al. Global time scale and regional stratigraphic reference scales of central and west europe, east europe, tethys, south china, and north america as used in the devonian–carboniferous–permian correlation chart 2003 (dcp 2003). Palaeogeography, Palaeoclimatology, Palaeoecology, 240(1-2):318–372, 2006.

Joseph E O’Reilly and Philip CJ Donoghue. The efficacy of consensus tree methods for summarizing phylogenetic relationships from a posterior sample of trees estimated from morphological data. Systematic biology, 67(2):354–362, 2017.

Joseph E O’Reilly and Philip CJ Donoghue. The effect of fossil sampling on the estimation of divergence times with the fossilised birth death process. Systematic biology, 2019. doi: 10.1093/sysbio/syz037.

Caroline Parins-Fukuchi. Use of continuous traits can improve morphological phylogenetics. Systematic Biology, 67(2):328–339, 2017.

Caroline Parins-Fukuchi. Bayesian placement of fossils on phylogenies using quantitative morphometric data. Evolution, 72(9):1801–1814, 2018.

John R Paterson, Gregory D Edgecombe, and Michael SY Lee. Trilobite evolutionary rates constrain the duration of the cambrian explosion. Proceedings of the National Academy of Sciences, 116 (10):4394–4399, 2019.

Matthew W Pennell, Jonathan M Eastman, Graham J Slater, Joseph W Brown, Josef C Uyeda, Richard G FitzJohn, Michael E Alfaro, and Luke J Harmon. geiger v2. 0: an expanded suite of methods for fitting macroevolutionary models to phylogenetic trees. Bioinformatics, 30(15): 2216–2218, 2014.

Mark N Puttick, Joseph E O’Reilly, Alastair R Tanner, James F Fleming, James Clark, Lucy Holloway, Jesus Lozano-Fernandez, Luke A Parry, James E Tarver, Davide Pisani, et al. Uncertain-tree: discriminating among competing approaches to the phylogenetic analysis of phenotype data. Proceedings of the Royal Society B: Biological Sciences, 284(1846):20162290, 2017.

Andrew Rambaut, Alexei J Drummond, Dong Xie, Guy Baele, and Marc A Suchard. Posterior Summarization in Bayesian Phylogenetics Using Tracer 1.7. Systematic Biology, 67(5):901–904, 04 2018. ISSN 1063-5157. doi: 10.1093/sysbio/syy032. URL https://dx.doi.org/10.1093/sysbio/syy032.

D.F. Robinson and L.R. Foulds. Comparison of phylogenetic trees. Mathematical Biosciences, 53 (1):131 – 147, 1981. ISSN 0025-5564. doi: https://doi.org/10.1016/0025-5564(81)90043-2. URL http://www.sciencedirect.com/science/article/pii/0025556481900432.

Fredrik Ronquist, Seraina Klopfstein, Lars Vilhelmsen, Susanne Schulmeister, Debra L. Murray, and Alexandr P. Rasnitsyn. A total-evidence approach to dating with fossils, applied to the early radiation of the hymenoptera. Systematic Biology, 2012. ISSN 10635157. doi: 10.1093/sysbio/sys058.

Klaus Peter Schliep. phangorn: phylogenetic analysis in r.Bioinformatics, 27(4):592–593, 2010.

Graham J. Slater. Iterative adaptive radiations of fossil canids show no evidence for diversity-dependent trait evolution. Proceedings of the National Academy of Sciences, 112(16):4897–4902, 2015. ISSN 0027-8424. doi: 10.1073/pnas.1403666111. URL https://www.pnas.org/content/112/16/4897.

Tanja Stadler. Sampling-through-time in birth–death trees. Journal of Theoretical Biology, 267(3): 396–404, 2010.

Tanja Stadler. Simulating trees with a fixed number of extant species. Systematic Biology, 60(5): 676–684, 2011. doi: 10.1093/sysbio/syr029. URL http://dx.doi.org/10.1093/sysbio/syr029.

Alycia L Stigall Rode. Systematic revision of the middle and late devonian brachiopods schizophoria (schizophoria) and ‘schuchertella’from north america. Journal of Systematic Palaeontology, 3(2): 133–167, 2005.

David F Wright. Bayesian estimation of fossil phylogenies and the evolution of early to middle paleozoic crinoids (echinodermata). Journal of Paleontology, 91(4):799–814, 2017a.

David F Wright. Phenotypic Innovation and Adaptive Constraints in the Evolutionary Radiation of Palaeozoic Crinoids. Scientific Reports, 7(1):13745, 2017b. ISSN 2045-2322. doi: 10.1038/s41598-017-13979-9. URL https://doi.org/10.1038/s41598-017-13979-9.

David F. Wright and Ursula Toom. New crinoids from the baltic region (estonia): fossil tip-dating phylogenetics constrains the origin and ordovician–silurian diversification of the flexibilia (echinodermata). Palaeontology, 60(6):893–910, 2017. doi: 10.1111/pala.12324. URL https://onlinelibrary.wiley.com/doi/abs/10.1111/pala.12324.

Chi Zhang, Tanja Stadler, Seraina Klopfstein, Tracy A Heath, and Fredrik Ronquist. Total-evidence dating under the fossilized birth–death process. Systematic Biology, 65(2):228–249, 2015.

