## Supplementary figures for "Ignoring fossil age uncertainty leads to inaccurate topology and divergence times in time calibrated tree inference"

### 1 Data used as basis for the simulation study

In order to select parameter values that would reflect the size and scale (in terms of taxon and character sampling, and time span) we first tallied 80 studies of Paleozoic invertebrate groups, which included trilobites (68%), brachiopods (18%) and crinoids (15%). The main features of these datasets are shown in Figure 1.

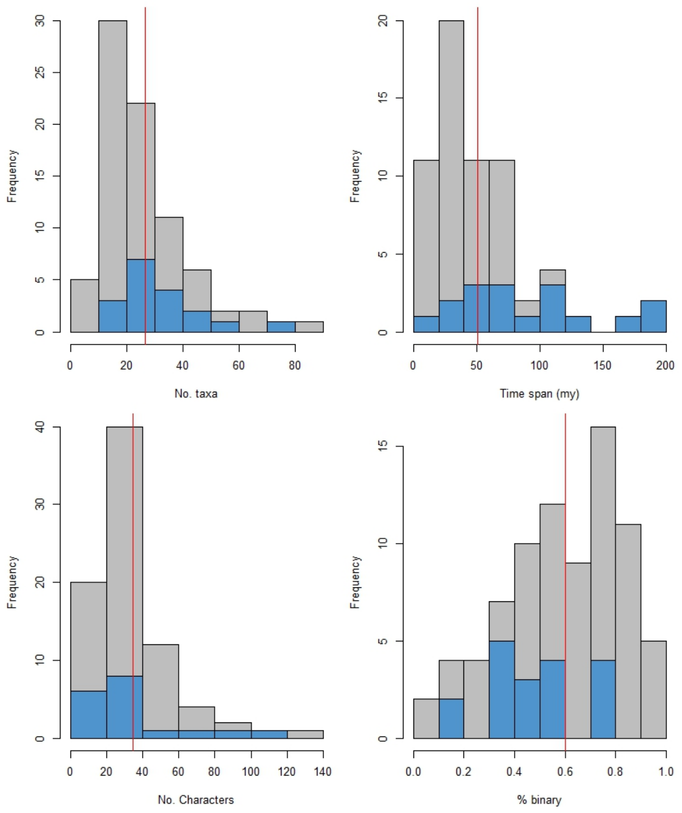


Figure 1: Histograms showing the features of datasets of Palaeozoic marine invertebrates. Species-level studies are shown in gray and the genus-level studies are in blue. The red line is the mean across everything.

### 2 Supplementary figures

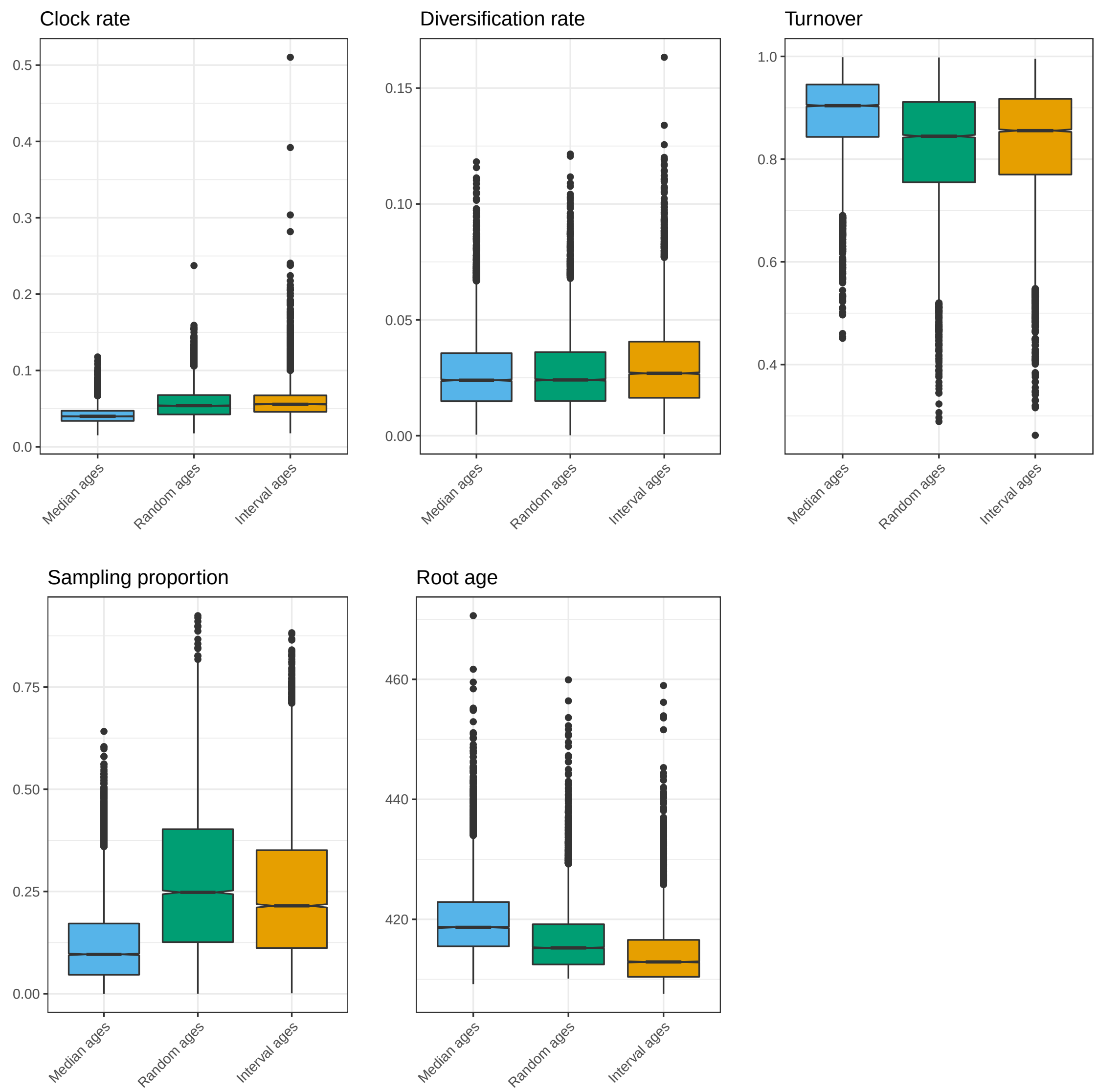


Figure 2: Parameter estimates obtained for the empirical brachiopods dataset, using different age handling methods.

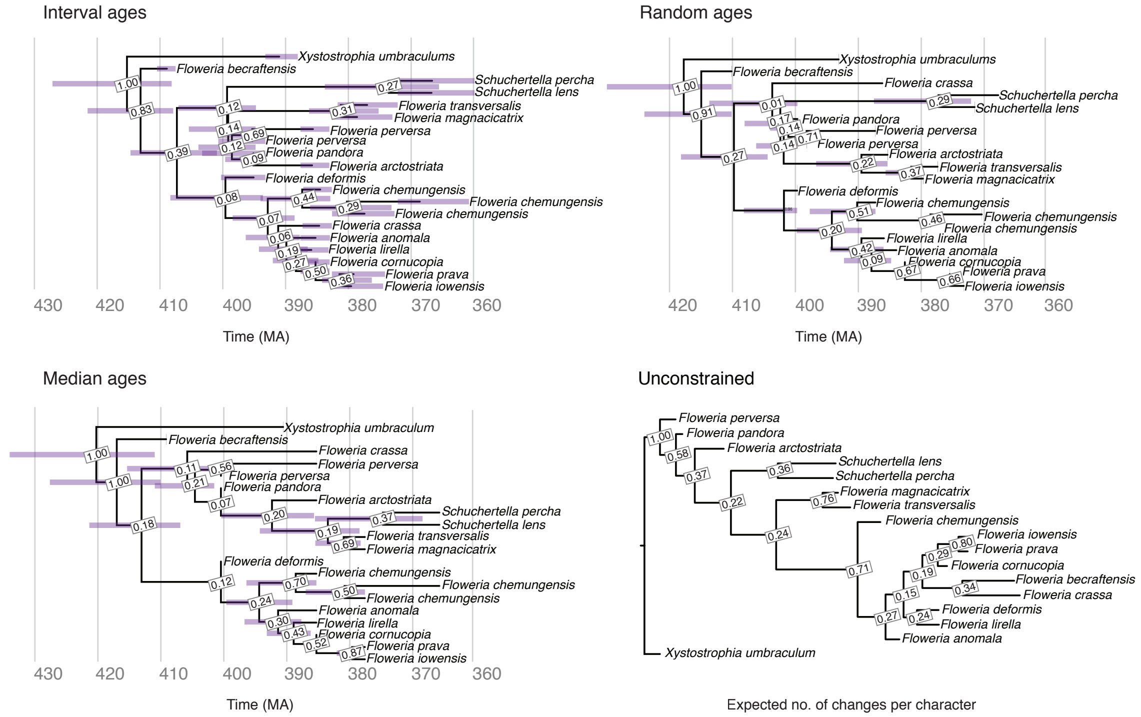


Figure 3: Brachiopod MCC trees obtained using the FBD analysis with interval ages, median ages and random ages, and unconstrained analysis. Posterior support is shown for each node for all trees. Error bars on the FBD trees show the 95% HPD interval for the age of each node, as well as the age of each fossil in the tree with interval ages.
